# A space-saving visual screening method, *Glycine max* FAST, for generating transgenic soybean

**DOI:** 10.1101/797282

**Authors:** Kosei Iwabuchi, Takashi L. Shimada, Tetsuya Yamada, Ikuko Hara-Nishimura

## Abstract

Soybean is an important crop plant for food and biofuel production, and there have been considerable efforts to develop transgenic soybean lines with higher seed oil contents and/or seed yields. However, the process of screening transgenic lines is laborious and requires a large amount of space. Here, we describe a powerful screening method, *Glycine max* Fluorescence-Accumulating Seed Technology (GmFAST), which is based on a seed-specific fluorescent marker. The marker is composed of a soybean seed-specific promoter coupled to the *OLE1-GFP* gene, which encodes GFP fused to the oil-body membrane protein OLEOSIN1 of *Arabidopsis thaliana*. We introduced the marker gene into cotyledonary nodes of *G. max* Kariyutaka via *Agrobacterium*-mediated transformation and regenerated heterozygous transgenic plants. OLE1-GFP-expressing soybean seeds can be selected nondestructively using a fluorescence stereomicroscope. Among T2 seeds, the most strongly fluorescent seeds were homozygous. GmFAST uses one-tenth of the growing space required for the conventional method. This space-saving method will contribute to facilitating transformation of soybean. OLE1-GFP was localized specifically to oil bodies in the cotyledon cells of seeds, but it did not affect oil content per seed, the size and density of the oil bodies, or oil composition. One of the homozygous lines (line #8) showed a 44% increase in the seed pod number, which resulted in 41% and 30% increases in seed yield and total oil production, respectively, compared with the wild type. In line #8, *OLE1-GFP* was inserted into the intron of Glyma13g30950, causing its overexpression. An increase in seed pod number was confirmed in *Arabidopsis thaliana* plants that overexpressed the Arabidopsis ortholog of Glyma13g30950, *E6L1*. These results suggest that line #8 is a valuable resource for agricultural and industrial applications. Taken together, GmFAST provides a space-saving visual and non-destructive screening method for soybean transformation, thereby increasing the chance of developing useful soybean lines.

## Introduction

Soybean (*Glycine max*) is an agriculturally important crop that produces oil-rich and protein-rich seeds (19% oil and 35%–40% protein) (Tidke et al. 2015). Therefore, soybean seeds are used as a source of food and nutraceuticals. Oil seeds have recently gained attention as biofuel resources with the increasing demand for energy in emerging countries. The total world production of soybean reached 276 million tons in 2013 and is increasing (FAOSTAT; http://faostat3.fao.org).

Soybean oil is composed of five fatty acids; palmitic acid (C16:0), stearic acid (18:0), oleic acid (18:1), linoleic acid (18:2), and linolenic acid (18:3)(Clemente and Cahoon 2009). These fatty acids are sequestered in a specific organelle, the oil body, which is 0.5 to 2.0 µm in diameter and surrounded by a lipid monolayer and membrane proteins including oleosin (Shimada and Hara-Nishimura 2010). Oleosin modulates the size of oil bodies (Abell et al. 1997; Shimada et al. 2008; Siloto et al. 2006) and the seed oil contents (Hu et al. 2009b; Shimada et al. 2008; Siloto et al. 2006). *Arabidopsis thaliana* has 16 oleosins, of which five (OLE1, OLE2, OLE3, OLE4, and OLE5) are expressed in seeds (Kim et al. 2002). Genetic engineering approaches based on oleosin function have been used to increase the seed oil content of rice; overexpression of a soybean oleosin increased the oil content of rice by 37%–46% compared with that of wild-type rice (Liu et al. 2013). However, whether this approach would also work in soybean seeds, where oil biosynthetic genes are reported to increase the oil content (Lardizabal et al. 2008; Rao and Hildebrand 2009), remains unknown.

The generation of transgenic plants involves two processes; introduction of the gene into plants, and screening of transgenic plants. Genes of interest are usually introduced via *Agrobacterium tumefaciens-*mediated transformation. The subsequent screening of transgenic plants is generally based on antibiotics or herbicides, since antibiotic- and herbicide-resistance genes are introduced into plants as selectable markers together with the gene(s) of interest. However, there are several problems associated with the screening process. Firstly, antibiotics/herbicides can inhibit plant growth. Secondly, screening requires aseptic techniques to prepare sterile seeds and agar plates containing antibiotics/herbicides, making the selection process time consuming and labor intensive.

To overcome these disadvantages, we previously established a method to select transgenic *Arabidopsis* and rice (*Oryza sativa*), designated as the FAST (Fluorescence-Accumulating-Seed Technology) method (Shimada et al. 2011; Shimada et al. 2010). This method relies on the FAST marker, in which a translational fusion gene encoding *Arabidopsis* OLE1 and green fluorescent protein (GFP), *OLE1-GFP*, is driven by a seed-specific promoter. The FAST marker is expressed in seeds, and the emission of GFP fluorescence in transgenic seeds allows for their easy and non-destructive isolation under a fluorescence stereomicroscope or blue LED handy-type instrument. Unlike conventional methods, the FAST method requires no aseptic techniques. This method reduced the time required to obtain homozygous transgenic plants from 7.5 months to 4 months in *Arabidopsis* (Shimada et al. 2010). The objective of the present study was to establish the FAST method in soybean. To this end, a GmFAST marker was generated, in which *OLE1-GFP* was expressed under the control of the *GLYCININ* promoter of *G. max* (Fischer and Goldberg 1982; Nielsen et al. 1989; Scallon et al. 1985). This screening system was used in combination with *Agrobacterium*-mediated transformation of *G. max* cotyledonary nodes (Yamada et al. 2010). Since the 11S globulin GLYCININ is highly expressed in seeds (Meinke et al. 1981; Schmidt et al. 2011), we examined the effect of OLE1-GFP on oil body organization and oil content in soybean seeds.

## Materials and Methods

### Plant materials and growth conditions of soybean plants

The Japanese soybean variety Kariyutaka was used as the wild type (WT), which was never placed in culture, and the OLE1-GFP-expressing plants were generated in the Kariyutaka background. Seeds were sown on compost and grown at 25°C under a 16-h light (white light at 100–150 µmol m^-2^ s^-1^), 8-h dark photoperiod. After germination, the position of plant pots relative to the light source was changed daily to ensure that all plants received the same amount of light.

### Plant materials and growth conditions of *Arabidopsis thaliana*

*Arabidopsis thaliana* ecotype Columbia (Col-0) was used as the wild type. Seeds, which had been surface-sterilized with 70% ethanol and dried, were sown on Murashige-Skoog (MS) agar plates (Wako) and grown at 22°C under continuous 100 µmol m^-2^ s^-1^white light for 2–3 weeks. Plants were then transferred to vermiculite and further grown at 22°C under the light.

### Plasmid DNA construct for the GmFAST-G marker

The *G. max* 11S globulin (GLYCININ) promoter (proGm11S, 1100 bp, accession AB353075) (Nakano et al. 2014) was cloned and a genomic fragment of proGm11S was amplified by PCR using the following primers: forward, 5′-GCAAGCTTGCGAATTCTCTCTTATAAAACACAAACAC-3′ and reverse, 5′-GCTCTAGAGCGTTTAAGGACCAATGGAGAGAATG-3′. Genomic DNA from *G. max* was used as the template for PCR. The amplified fragment was purified and amplified with Taq polymerase with the addition of adenine. The PCR product was inserted into the pGEM-T-Easy (Promega, Madison, WI, USA) vector to produce pGEM-proGm11S. Then, pGEM-proGm11S was treated with *Hind*III and *Xba*I to cut out the proGm11S fragment. pMDC123-IG (Yamada et al. 2010) was also treated with *Hind*III and *Xba*I to remove the 35S promoter region. The two fragments were ligated to produce the pMDC123-Gm11SIG vector.

A DNA fragment containing proGm11S, the *OLE1-GFP* gene, and the 35S terminator (ter35S) was amplified by overlapping PCR. In the first PCR, the proGm11S fragment was amplified using the following primers: forward, 5′-CACCGAATTCTCTCTTATAAAACACAA-3′ and reverse, 5′-TGTATCCGCCATGTTTAAGGACCAATGGAG-3′. The template for PCR was pMDC123-Gm11SIG. The *OLE1-GFP*::ter35S fragment was amplified using the following primers: forward, 5′-TGGTCCTTAAACATGGCGGATACAGCTAGA-3′ and reverse, 5′-GAATTCCGACGTCGCATGCCTGCAGGTCA-3′. The template for PCR was pBGWF7-proOLE1::OLE1-GFP (Karimi et al. 2002; Shimada et al. 2010). In the second PCR, the proGm11S::*OLE1-GFP*::ter35S fragment was amplified using the following primers: forward, 5′-CACCGAATTCTCTCTTATAAAACACAA-3′ and reverse, 5′-GAATTCCGACGTCGCATGCCTGCAGGTCA-3′. The first PCR products were used as the templates for PCR. The proGm11S::*OLE1-GFP*::ter35S fragment was inserted into pENTER/D-TOPO (Invitrogen, Carlsbad, CA, USA) by the TOPO reaction to produce pENTR/proGm11S::*OLE1-GFP*::ter35S. The proGm11S::*OLE1-GFP*::ter35S fragment was amplified for use in the In-Fusion cloning system (Clontech Laboratories, Mountain View, CA, USA) using the following primers: forward, 5′-CCAAGCTTGCGAATTCTCTCTTATAAAACACAAACACAATT-3′ and reverse, 5′-CCATGATTACGAATTCCGACGTCGCATGCCTGCAGGTCACT-3′. The template for PCR was pENTR/proGm11S::*OLE1-GFP*::ter35S. Finally, the fragment was inserted into the *Eco*RI site of pMDC123 (Curtis and Grossniklaus 2003) using the In-Fusion cloning system to produce the expression vector GmFAST-G.

### Plasmid DNA construct for *E6L1* overexpression

The full-length genomic fragment of *A. thaliana E6L1*, starting from nucleotide 1 (corresponding to A of the start codon ATG) to nucleotide 804, was amplified by PCR using the primer set: 5’-GCAGGCTCCGCGGCCATGGCTTTTTCCACTAGCTC-3’ (forward) and 5’-AGCTGGGTCGGCGCGTCATGGAGTGATCTGATCTC-3’ (reverse). The PCR product was subcloned with In-Fusion HD Cloning Kit (Takara Bio Inc.) into pENTR/D-TOPO (Invitrogen) which was digested by *Not*I and *Asc*I. The subcloned sequence was then transferred into the pH2GW7 vector (Karimi et al. 2002) via a recombination reaction with LR Clonase (Invitrogen), yielding the pH2GW7-E6L1 vector.

### Transformation of soybean plants

*Agrobacterium*-mediated transformation was performed as described previously (Yamada et al. 2010). The cotyledonary nodes of Kariyutaka were prepared for infection with *A. tumefaciens* EHA105 harboring GmFAST-G. The T0 plants were selected based on resistance to glufosinate-ammonium (Sigma-Aldrich, St. Louis, MO, USA). The inheritance of the transgene was confirmed based on seedling resistance to Basta (Bayer Crop Science, Monheim, Germany). In addition, mature seeds emitting GFP were selected under a fluorescence stereomicroscope (MVX10, Olympus, Tokyo, Japan). Three independent lines (#30, #2, and #8) were established, and homozygous T3 plants were used in further analyses.

### Stable transformation of Arabidopsis plants

The binary vector pH2GW7-E6L1 was introduced into *A. thaliana* (Col-0) plants with *Agrobacterium tumefaciens* (strain GV3101) using the floral dip method (Bechtold and Pelletier 1998) to generate *E6L1* overexpression (E6L1-OX) plants.

### SDS-PAGE and immunoblotting for soybean

Three dry seeds were broken into small pieces with a hammer and ground to a powder with a mortar and pestle in liquid nitrogen. Fifty milligrams of each powdered sample was diluted in 10 ml sample buffer [50 mM Tris-HCl, pH 6.8, 2% w/v sodium dodecyl sulfate (SDS), 6% v/v β-mercaptoethanol, and 10% v/v glycerol], mixed well, boiled at 99°C for 5 min, and then centrifuged at 9,000 × *g* for 1 min. A 1-mL aliquot of the supernatant was mixed with 50 µl bromophenol blue, and then the extracted proteins were separated by SDS-PAGE on a precast gel (Bio-Rad, Hercules, CA, USA) at a constant voltage of 200 V for 45 min. The gel was stained with Coomassie Brilliant Blue (CBB).

For immunoreaction experiments, gel-separated proteins were transferred onto a polyvinylidene fluoride membrane using the iBlot system (Invitrogen, Carlsbad, CA, USA). The membrane was blocked with skim milk in TBST buffer (20 mM Tris-HCl, pH 7.5, 150 mM NaCl, and 0.05% Tween-20). The primary antibody reaction was performed with anti-OLE1 diluted 1:2000 or anti-GFP diluted 1:5000 in TBST buffer for 1 h. The secondary antibody reaction was performed with anti-mouse IgG-HRP diluted 1:5000 or anti-rabbit IgG-HRP diluted 1:5000 for 30 min. After the reaction of HRP-conjugated secondary antibody with ECL (GE Healthcare Life Sciences, Buckinghamshire, United Kingdom), chemiluminescence was visualized using a LAS-3000 LuminoImage analyzer (FujiFilm, Tokyo, Japan).

### DNA isolation and PCR analysis for soybean

Dry seeds were broken into small pieces with a hammer and then ground to a powder with a mortar and pestle in liquid nitrogen. Genomic DNA was purified from seeds of *G. max* with a DNeasy plant mini kit (Qiagen, Hilden, Germany) as per the manufacturer’s instructions. The primer sets used to amplify the OLE1, GFP, and OLE1-GFP fragments were as follows: 5′-ACCCACAGGGATCAGACAAG-3′ (OLE1 forward) and 5′-GTTCCCCACCAGTATGTTGC-3′ (OLE1 reverse), and 5′-CACCATGGTGAGCAAGGGCGAGGAGCTGTT-3′ (GFP forward) and 5′-TCAGAGATCTCCCTTGTACAGCTCGTCCAT-3′ (GFP reverse). The PCRs were performed with MightyAmp DNA polymerase (Takara, Otsu, Japan) under the following conditions: 40 cycles of 98°C for 10 s, 55°C for 30 s, and 72°C for 60 s.

### Fluorescence microscopy for soybean

Dry seeds were fixed in fixation buffer (50 mM PIPES, 10 mM EGTA, and 5 mM MgSO_4_, pH 7.0) containing 2% formaldehyde and 0.3% glutaraldehyde overnight. The GFP fluorescence of seed cotyledons was observed under a fluorescence stereomicroscope (Zeiss, Oberkochen, Germany) or a confocal laser scanning microscope (LSM780 META; Zeiss). The mean GFP fluorescence level of dry seeds was determined using Image J software (http://rsb.info.nih.gov/ij).

To visualize lipids, the fixed seed cotyledons were stained with 5 µg/ml Nile red and observed under a confocal laser scanning microscope (Zeiss LSM780 META).

### Transmission electron microscopy for soybean

Dry seeds were cut into approximately 1 × 1 mm pieces using a razor blade and fixed with 4% paraformaldehyde and 2% glutaraldehyde in 0.05 M cacodylate buffer, and then with 2% osmium tetroxide in 0.05 M cacodylate buffer. After dehydration, samples were embedded in Quetol 651 resin (Electron Microscopy Sciences, Hatfield, PA, USA). Ultrathin sections (80 nm) were prepared, stained with uranyl acetate in lead stain solution, and observed under a transmission electron microscope (JEM-1400Plus; JEOL, Tokyo, Japan).

### Number of seeds and seed pods, and seed weight measurements for soybean

The T3 plants were grown under the conditions described above until the seed pods were completely dry, and then the seeds were harvested together with their pods. The number of seeds per plant was counted and the total seed weight per plant was measured with a digital electronic balance (XS105DU; Mettler Toledo, Greifensee, Switzerland). The number of pods per plant was also counted, and the percentage of pods with one, two, or three seeds on each plant was calculated.

### Fatty acid content analysis for soybean

The fatty acid content of T4 dry seeds was analyzed as described previously (Yamada et al. 2014). Mature seeds were ground using a mortar and pestle. The fatty acid content (palmitic, oleic, linoleic, and linolenic acids) was determined by gas chromatography based on the ratio of the areas of respective peaks to that of heptadecanoic acid. Lipid content was calculated as the total fatty acid content.

### An adapter ligation-mediated PCR for soybean

The site of *OLE1-GFP* insertion in the genome of OLE1-GFP transgenic line #8 was determined by an adapter ligation-mediated PCR as reported previously (O’Malley et al. 2007). Briefly, total DNA was isolated from seeds with a DNeasy Plant Mini Kit (Qiagen) according to the manufacturer’s instructions. Total DNA was digested with restriction enzymes (Hind III and Eco RI), and the digested DNA was ligated to the following primers: 5’-GTAATACGACTCACTATAGGGCACGCGTGGTCGACGGCCCGGGCTGC-3’ (long strand of adapter 1 for Hind and Eco), 5’-AGCTGCAGCCCG-3’ (short strand of adapter Hind), and 5’-AATTGCAGCCCG-3’ (short strand of adapter Eco). The DNA was amplified by a nested PCR using the following primer sets: 5’-AGTCGACCGTGTACGTCTCC-3’ (pMDC123_LB1) and 5’-GTAATACGACTCACTATAGGGC-3’ (AP1) for 1^st^ PCR; 5’-GTTTCTGGCAGCTGGACTTC-3’ (pMDC123_LB2) and 5’-TGGTCGACGGCCCGGGCTGC -3’ (AP2) for 2^nd^ PCR, 5’-AAAAACGTCCGCAATGTGTT-3’ (pMDC123_LB3) and 5’-TGGTCGACGGCCCGGGCTGC -3’ (AP2) for 3^rd^ PCR. The 3^rd^ PCR products were subjected to agarose gel electrophoresis, and DNA fragments of interest were excised from the gel and purified. The DNA fragments were sequenced using a genetic analyzer (3130xl; ABI). Insertion of OLE1-GFP into Glyma13g30950 was confirmed by genotyping PCR using the following primer sets: 5’-ACTGTAGGCTTGATGCCACT-3’ (forward) and 5’-TCCTCTACGTCATTCGATGG-3’ (reverse); 5’-AAAAACGTCCGCAATGTGTT-3’ (pMDC123_LB3) and 5’-TCCTCTACGTCATTCGATGG-3’ (reverse).

### Reverse transcription PCR for soybean

Total RNA was isolated from soybean leaves using RNeasy Plant Mini Kit (Qiagen) according to the manufacturer’s instructions. Total RNA was subjected to first-strand cDNA synthesis using Ready-To-Go RT-PCR Beads (GE Healthcare), and the cDNA was amplified by PCR using the following primer sets: 5’-TTTTCCTCACCACCCTTTTG -3’ (forward) and 5’-CGGAATGAAAGGTGGTTGTT-3’ (reverse); 5’-ATCTTGACTGAGCGTGGTTATTCC-3’ (forward) and 5’-GCTGGTCCTGGCTGTCTCC-3’ (reverse) for *ACTIN11*(Hu et al. 2009a). The size of PCR products of Glyma13g30950 and *ACTIN11* was 164 and 126 bp, respectively.

### Reverse transcription PCR for Arabidopsis

Total RNA was isolated from Arabidopsis seedlings using RNeasy Plant Mini Kit (Qiagen). The RNA was used for cDNA synthesis using ReverTra Ace qPCR RT Master Mix (Toyobo), followed by PCR using the following primer sets: 5’-ATGGCTTTTTCCACTAGCTC-3’ (forward) and 5’-TCATGGAGTGATCTGATCTC-3’ (reverse) for *E6L1*, and 5’-ACTGGAGGTTTTGAGGCTGGTAT-3’ (forward) and 5’-GCACCGTTCCAATACCACCAATC-3’ (reverse) for *EF1a*. The size of PCR products of *E6L1* and *EF1a* was 804 and 494 bp, respectively.

### Number of seeds and seed pods, and seed weight measurements for Arabidopsis

Wild-type and E6L1-OX plants (T2 generation) were grown until the seed pods were completely dried. For confirming whether the T2 plants contained the transgene, the following primer sets were used for genotyping by PCR: 5’-ATGAAAAAGCCTGAACTCACCGC-3’ (forward) and 5’-CTATTCCTTTGCCCTCGGACGAG-3’ (reverse) for hygromycin-resistant gene. The seed pods in each plant were counted before seed harvesting. The seeds were harvested in each plant, and then the total seed weight per plant was measured with a digital electronic balance (GH-120; A&D Company, Tokyo, Japan). The weight of 100 seeds were measured with the digital electronic balance (GH-120) for calculating the weight of one seed. According to the data, the number of seeds per plant and per pod were calculated.

## Results and Discussion

### Identification of transgenic soybean seeds by GmFAST

The FAST system for soybean was established by constructing the GmFAST-G vector, which contains the *OLE1-GFP* fusion gene driven by the *G. max* 11S globulin promoter (Fig. 1A). The GmFAST-G vector was introduced into cotyledonary nodes of *G. max* Kariyutaka through *Agrobacterium*-mediated transformation. After the regeneration of transgenic plants from independent cotyledonary nodes, T1 mature seeds were harvested and grown to obtain the T2 seed population, which contained homozygous, heterozygous, and WT seeds. Next, we carried out T2 seed selection based on the GmFAST method. The GFP fluorescence levels are reflective of the GmFAST-G vector copy numbers: two and one in homozygotes and heterozygotes, respectively (Shimada et al. 2010). Therefore, we selected strongly fluorescent seeds from independent T2 seed populations using a fluorescence stereomicroscope and finally obtained T3 seeds from three independent transgenic lines designated as #2, #8, and #30 (Fig. 1B). The GFP fluorescence intensity levels of lines #2 and #8 were twice that of WT, whereas the GFP fluorescence of line #30 was lower and similar to that of WT (Fig. 1C). The homozygosity of these lines was confirmed by a non-destructive segregation analysis of the T3 seeds using a fluorescence stereomicroscope. As expected, all the T3 seeds from each line emitted GFP fluorescence, indicating that they were homozygous. The insertion of the *OLE1-GFP* gene in all three transgenic lines was also confirmed by genotyping (Fig. 1D, upper). An immunoblot analysis showed that the OLE1-GFP protein was expressed in lines #2 and #8, but was not expressed at detectable levels in line #30 (Fig. 1D, middle). Thus, the GmFAST method enables the isolation of homozygous plants in the T2 generation.

**Fig. 1.**
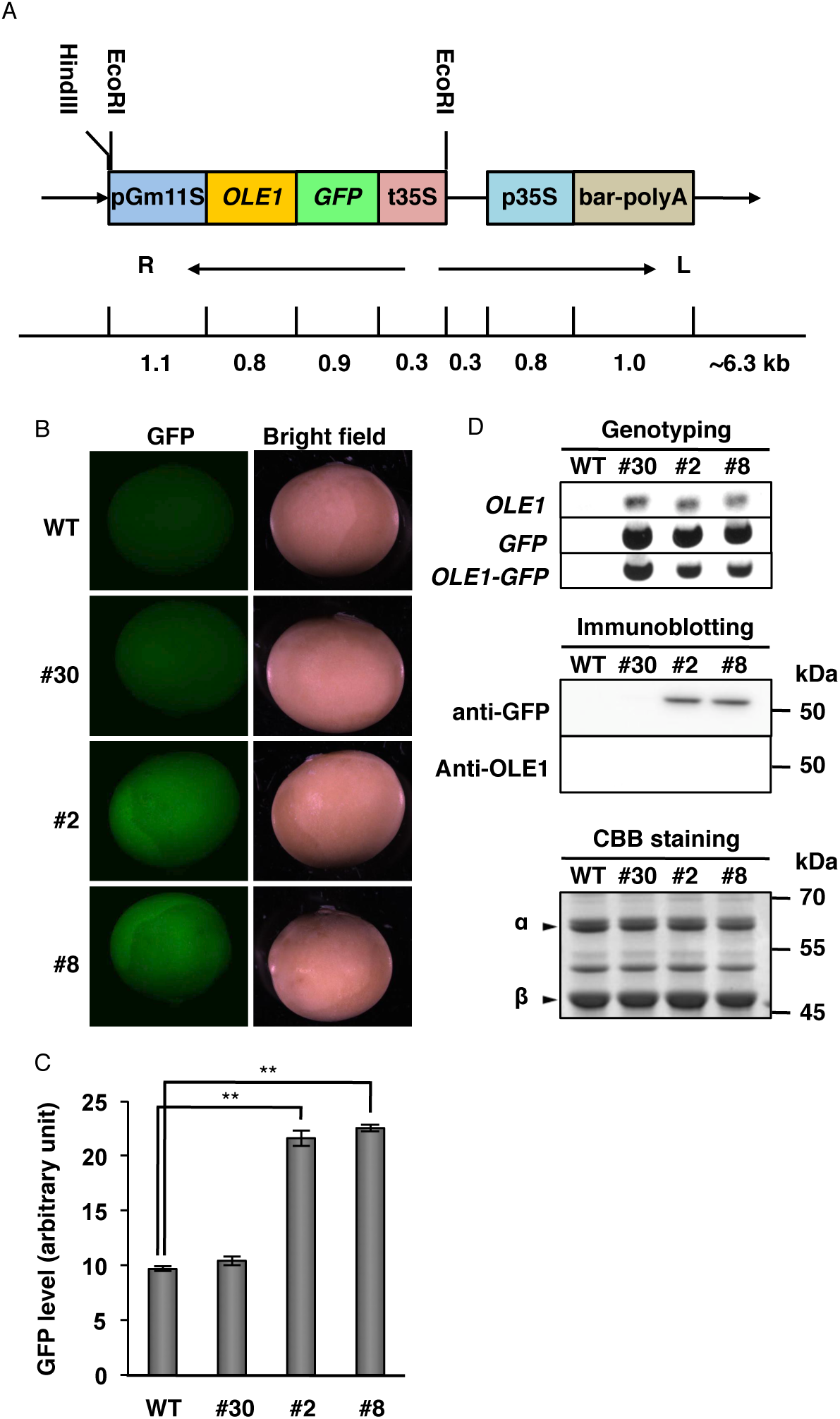
Expression of OLE1-GFP in transgenic soybean seeds. (A) Schematic diagram of the GmFAST-G vector. *Arabidopsis OLE1* was fused to *GFP* and expressed under the control of the soybean 11S globulin promoter. (B) Representative GFP fluorescence images of dry mature seeds of wild type (WT) and OLE1-GFP transgenic lines (#2, #8, and #30). (C) Quantification of GFP fluorescence determined using data from (B). The mean fluorescence of 10 seeds was determined for each plant. Data represent means ± SEMs (*n* = 6 plants; \*\**P* < 0.01, Student’s *t* test). (D) Expression of OLE1-GFP in dry seeds of WT and OLE1-GFP transgenic lines (#2, #8, and #30). Genotyping was performed with specific primers for *OLE1, GFP*, and *OLE1-GFP* (upper), and immunoblotting was performed with anti-GFP and anti-OLE1 (middle). CBB staining was conducted as a loading control for immunoblotting (lower). Arrowheads indicate α and β subunits of 7S globulin.

### Intracellular localization of OLE1-GFP in transgenic soybean seeds

An analysis of the localization of OLE1-GFP in transgenic seeds showed that it was exclusively expressed in seed cotyledons in lines #2 and #8 (Fig. 2A; #2 and #8). In the palisade mesophyll cells of cotyledons, OLE1-GFP showed a meshwork-like expression pattern (Fig. 2B, left), similar to the distribution pattern of the oil bodies stained with Nile red (Fig. 2B, right). These results suggested that OLE1-GFP was functional in oil bodies.

**Fig. 2.**
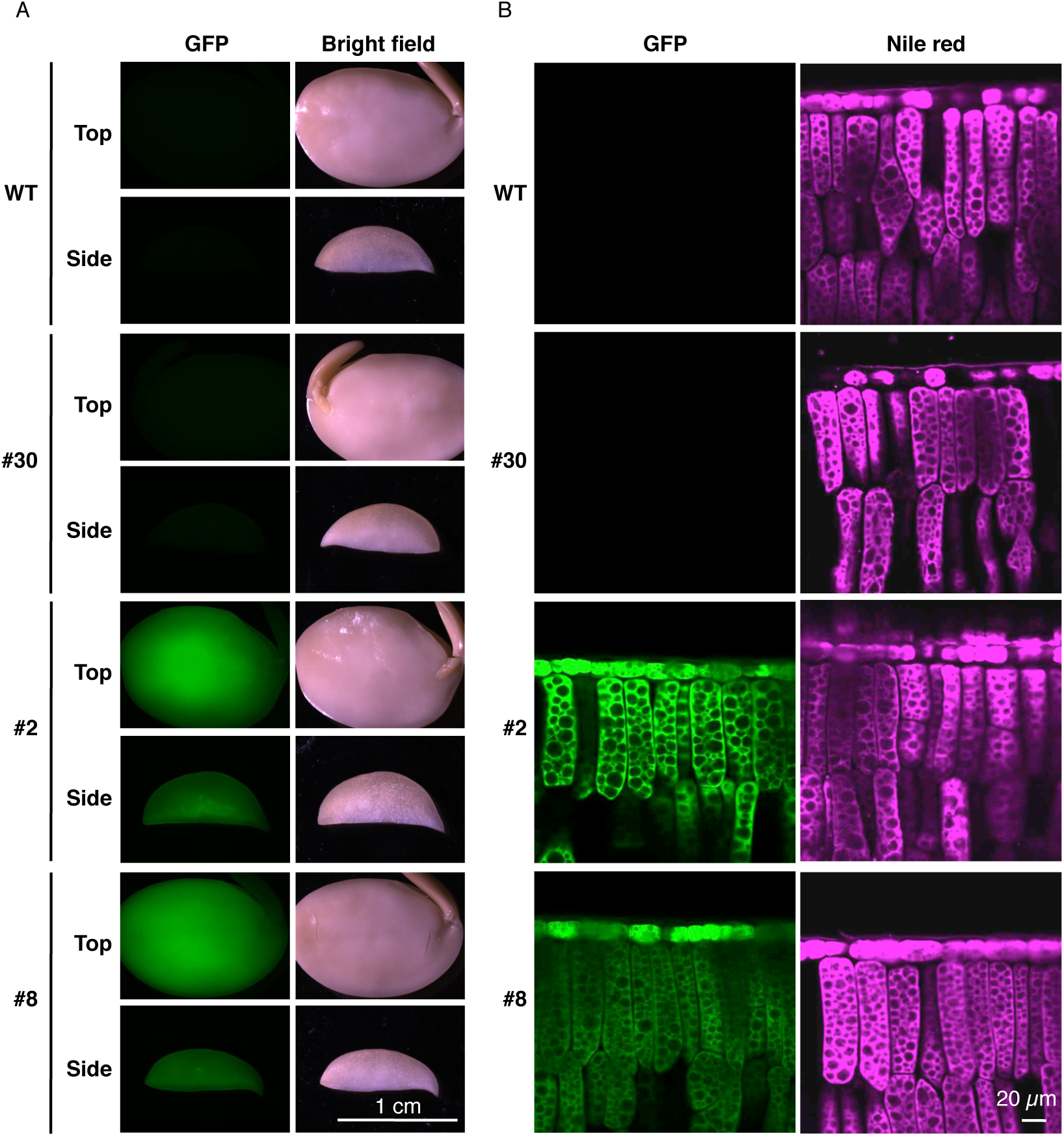
Localization of OLE1-GFP in transgenic soybean seeds. (A) Representative images of seed cotyledons of wild type (WT) and OLE1-GFP lines (#2, #8, and #30) after fixation. Top (upper panels) and side (lower panels) views are shown. (B) Side-view images of adaxial epidermal and palisade mesophyll cells of cotyledons in WT and OLE1-GFP transgenic lines (#2, #8, and #30), showing the intracellular localization of OLE1-GFP (left panels) and lipid droplets using Nile red staining (right panels).

Next, we examined the effects of OLE1-GFP on oil body organization in transgenic seeds using transmission electron microscopy. As previously reported (Herman and Larkins 1999; Schmidt et al. 2011; Yamada et al. 2014), seed cotyledon cells contained many oil bodies and several protein storage vacuoles (PSVs) in all the plants examined (Fig. 3, left). No differences in the size and density of oil bodies were observed between WT seeds and seeds of lines #30 and #2 (Fig. 3, right). This could be attributed to the low expression levels of OLE1-GFP. However, compared with WT seeds, line #8 seeds had oil bodies that were relatively small and densely distributed (Fig. 3, right). Line #8 also exhibited an altered PSV organization (Fig. 3, left). These results were consistent with previous reports that the oleosin contents affects both oil-body sizes and PSV organization in *Arabidopsis* and soybean (Schmidt and Herman 2008; Shimada et al. 2008; Siloto et al. 2006).

**Fig. 3.**
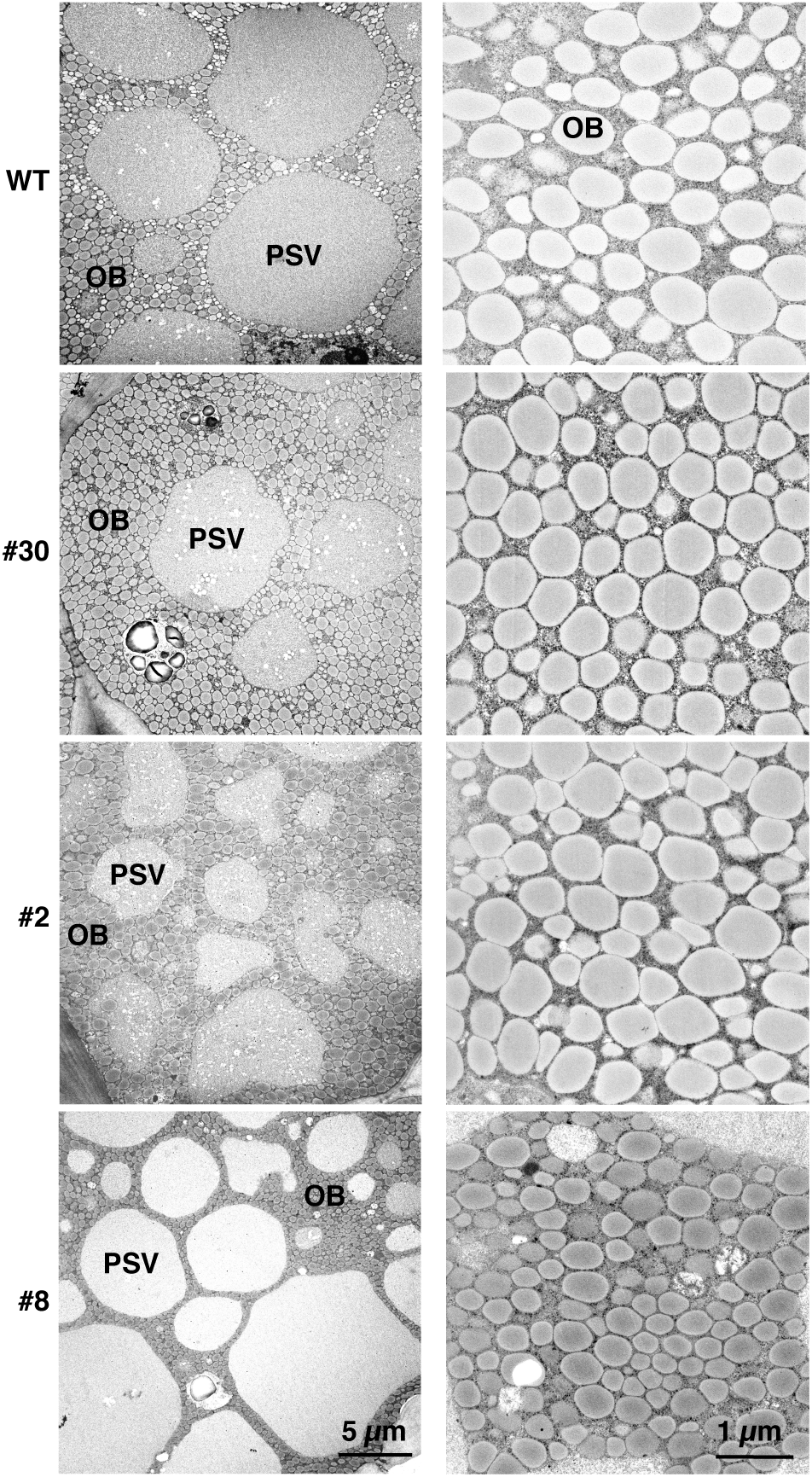
Morphology of protein storage vacuoles and oil bodies in transgenic soybean seeds. Representative transmission electron micrographs of mature seeds in wild type (WT) and OLE1-GFP transgenic lines (#2, #8, and #30) showing protein storage vacuoles (PSVs) and oil bodies (OBs). High-magnification images of OBs are shown in the right panels.

### Seed production characteristics and seed oil contents in transgenic soybean plants

The seed production of each transgenic line was examined. Under our experimental conditions, line #8 showed a 44% increase in seed pod number per plant compared with WT (Fig. 4A). Line #8 also showed 41% increases in total seed weight per plant (Fig. 4B) and 47% seed grain number per plant compared with WT (Fig. 4C). However, the average weight per seed (Fig. 4D) and the proportions of one-, two-, and three-seed pods (Fig. 4E) did not differ significantly between WT and line #8. Lines #2 and #30 did not show any changes in the above parameters compared with WT (Fig. 4). Considering the comparable expression levels of OLE1-GFP in lines #2 and #8 (Fig. 1C), the higher seed production of #8 might not be related to OLE1-GFP function. Instead, the insertion position of the *OLE1-GFP* gene in the genome might be responsible for the higher seed production in line #8.

**Fig. 4.**
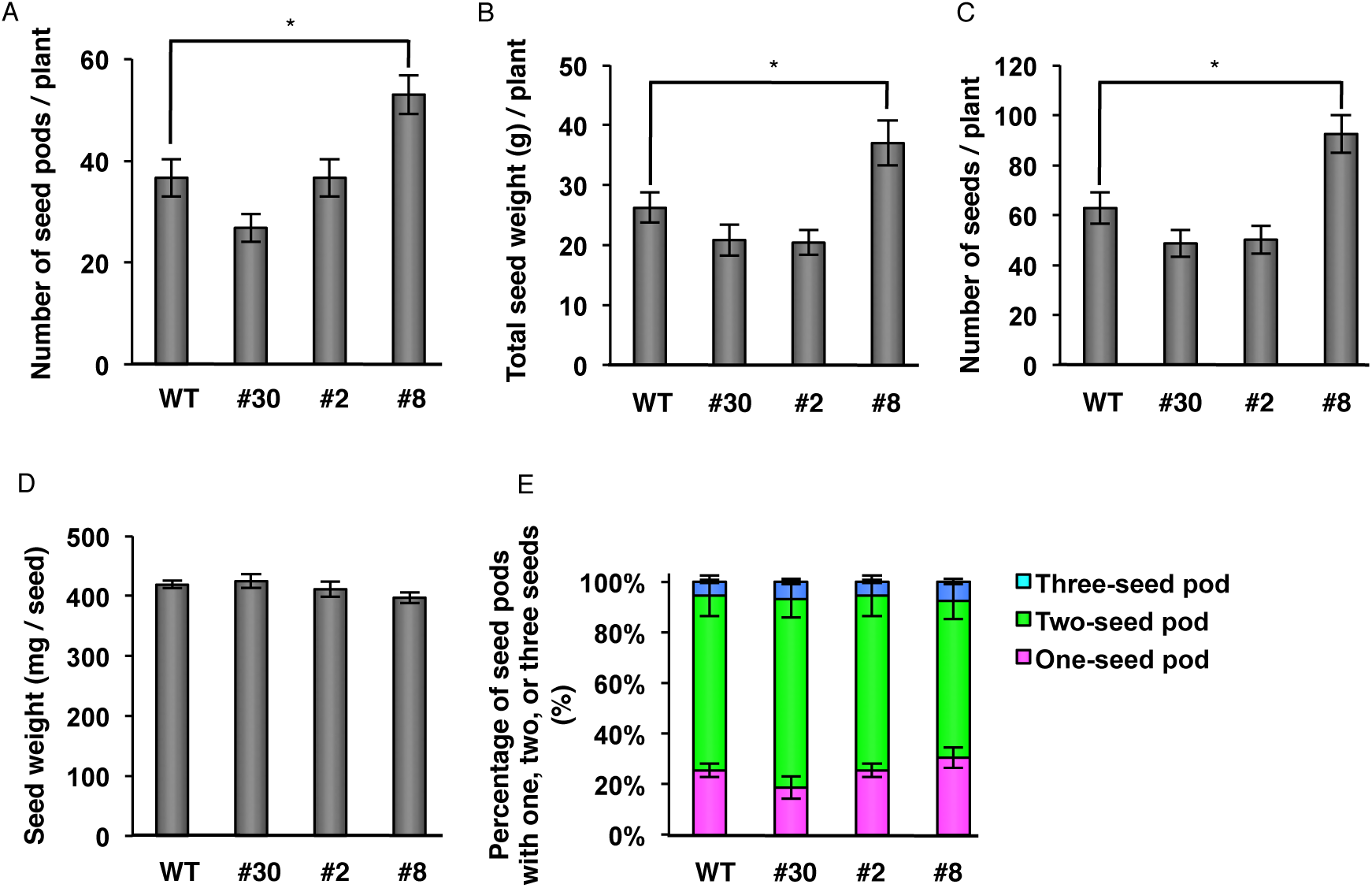
Seed production in transgenic soybean plants. Dry mature seeds were harvested from wild type (WT) and OLE1-GFP lines (#2, #8, and #30). (A) Number of seed pods per plant. (B) Total weight of seeds per plant. (C) Number of seeds per plant. (D) Weight per seed. (E) Percentages of pods with one, two, or three seeds. Data are means ± SEMs (*n* = 6 plants; * *P* < 0.05, Student’s *t* test).

Next, we examined the fatty acid contents of transgenic seeds. Line #8 showed a 30% increase in total oil production per plant compared with WT. The amounts of palmitic acid (C16:0), stearic acid (C18:0), and linolenic acid (C18:3) were higher in seeds of line #8 than in seeds of WT (Fig. 5A). However, the total fatty acid content based on seed weight was similar among all three transgenic lines (including line #8) and WT (Fig. 5B). Consequently, line #8 may be a valuable resource for agricultural and industrial applications.

**Fig. 5.**
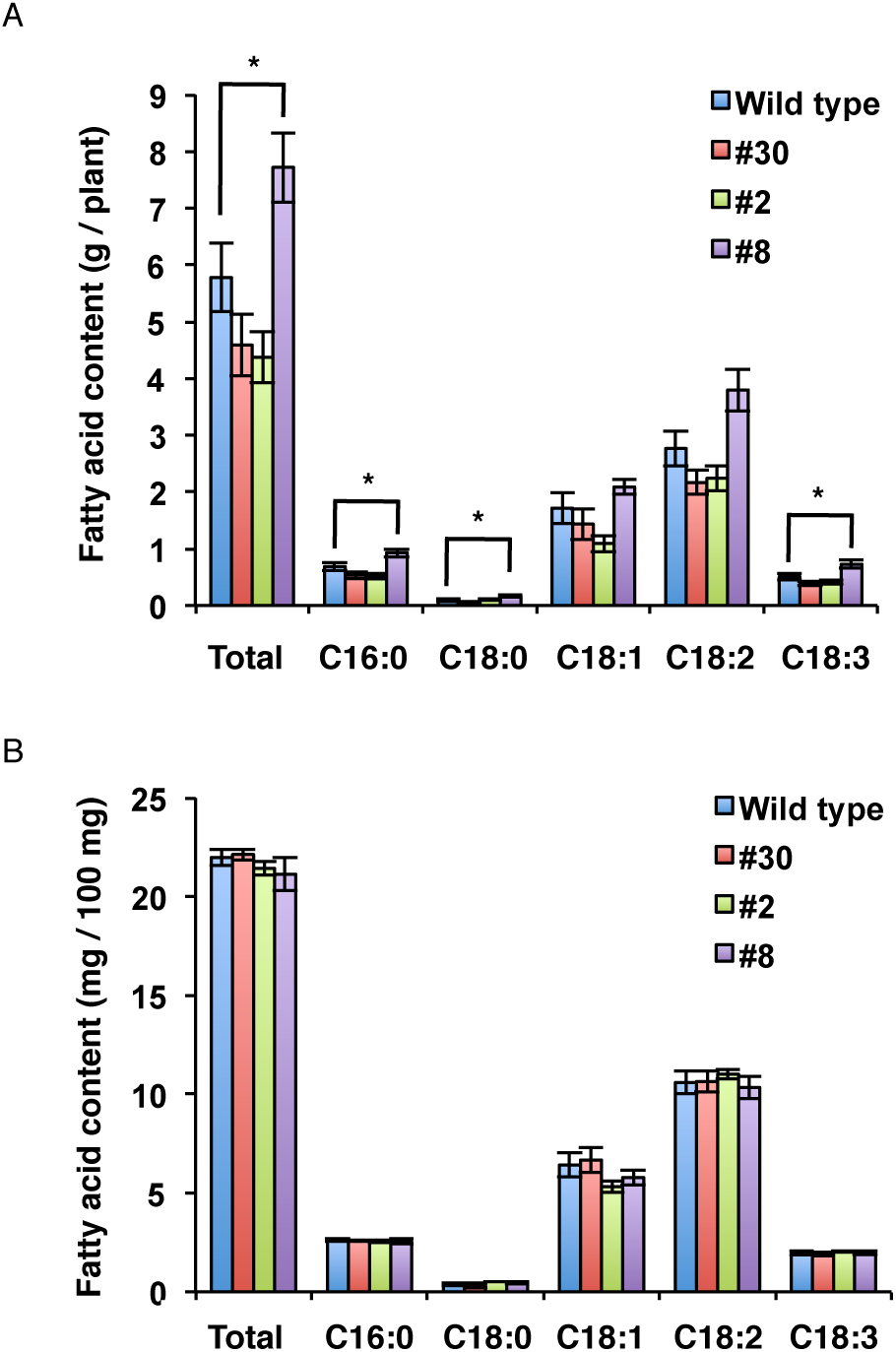
Fatty acid composition in transgenic soybean seeds. The fatty acid contents of total seeds (A) and a 100 mg of seeds (B) from the wild type and OLE1-GFP transgenic lines (#2, #8, and #30). C16:0, palmitic acid; C18:0, stearic acid; C18:1, oleic acid; C18:2, linoleic acid; C18:3, linolenic acid. Data represent means ± SEMs (*n* = 6 plants; \**P* < 0.05, Student’s *t* test).

In a previous study, the oil content increased by ∼40% in the transgenic rice grains that expressed soybean oleosin under the control of the RICE EMBRYO GLOBULIN-2 PROTEIN (REG-2) promoter (Liu et al. 2013). However, no increase in the oil content based on seed weight was observed in soybean seeds that expressed the oleosin gene *OLE1* under the control of the soybean 11S globulin (GLYCININ) promoter (Fig. 5B), possibly because soybean seeds inherently have much higher oil contents than rice grains.

### Site of the *OLE1-GFP* insertion in the transgenic soybean line #8 genome

Because the phenotype of line #8 could be result from an insertion mutation of *OLE1-GFP*, we performed an adapter ligation-mediated PCR (O’Malley et al. 2007) to locate the insertion site of *OLE1-GFP* in its genome. *OLE1-GFP* was inserted in the intron of Glyma13g30950, and this was confirmed using genotyping PCR (Fig. 6A and 6B). Intriguingly, the OLE-GFP insertion enhanced the transcript level of Glyma13g30950 in #8, compared with the wild type, #2, and #30 (Fig. 6C).

**Fig. 6.**
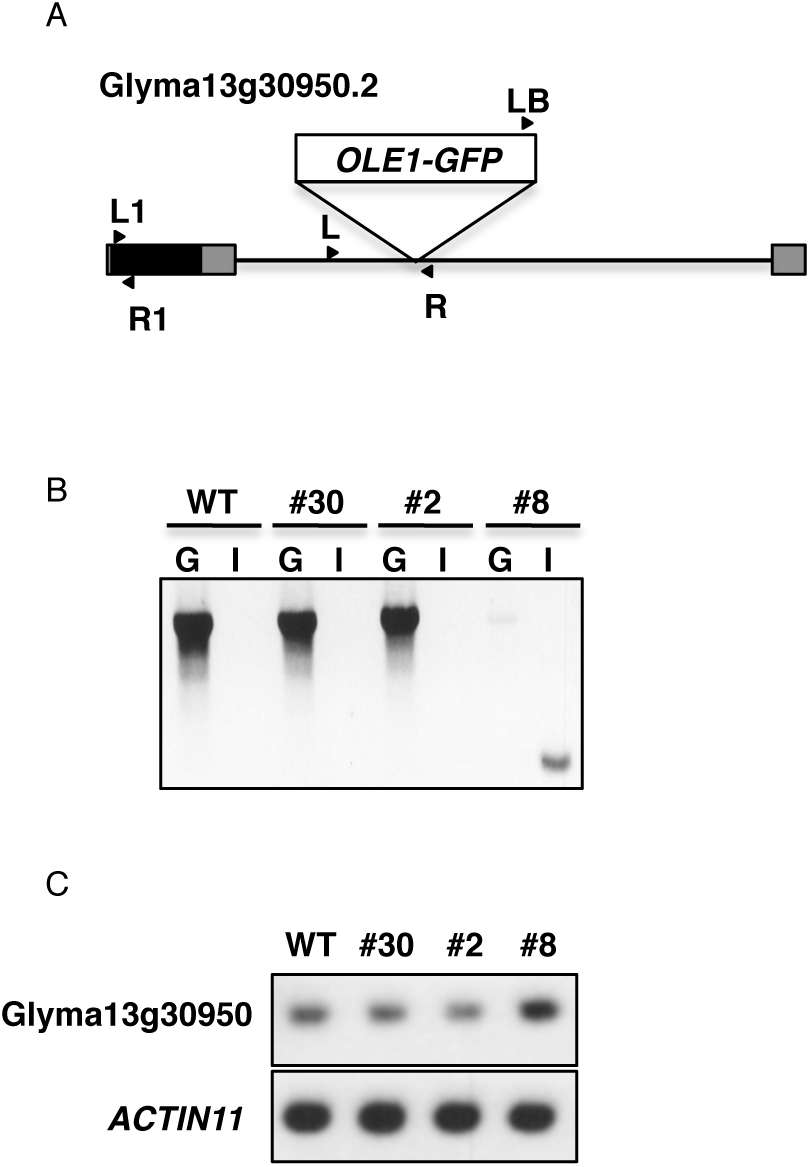
Site of the *OLE1-GFP* insertion in the transgenic soybean line #8 genome. (A) Schematic representation of Glyma13g30950.2, showing the site of the *OLE1-GFP* insertion. Gray boxes, untranslated regions; black box, exon; solid line, intron; arrow heads, primer positions used in B and C. (B) Genotyping in the wild type (WT) and OLE1-GFP-expressing lines, #2, #8, and #30. G, genomic fragments amplified with a pair of forward and reverse Glyma13g30950-specific primers (L and R in A); I, an inserted fragment amplified with a left border primer for pDMC123 vector (LB in A) and Glyma13g30950 reverse primer (R in A). (C) RT-PCR of Glyma13g30950 and *ACTIN11* (control) transcripts in leaves of the WT and OLE1-GFP-expressing lines, #2, #8, and #30. Fragments were amplified with Glyma13g30950-specific primers (L1 and R1 in A).

### The effects of E6L1 on seed pod number in Arabidopsis

To support our hypothesis that Glyma13g30950 is involved in seed pod formation, we investigated the function of the Glyma13g30950 ortholog in Arabidopsis. From a BLAST analysis using the amino acid sequence of Glyma13g30950.2, a candidate for an Arabidopsis orthologs, an *E6-like1* gene (*E6L1*; At2g33850), was found. Then, we generated transgenic Arabidopsis plants overexpressing *E6L1* (E6L1-OX) (Fig. S1). Of three lines analyzed (#2, #6, and #12), line #2 exhibited a statistically significant increase in seed pod number compared with WT (Fig. 7A), although there were no remarkable changes between WT and the three E6L1-OX lines in the total seed weight per plant, total seed grain number per plant, the weight per seed, or the seed number per pod (Fig. 7B–E). These results suggest that E6L1 is involved in regulating the seed pod number. According to the Arabidopsis eFP browser (http://bar.utoronto.ca/efp/cgi-bin/efpWeb.cgi), *E6L1* is expressed weakly at the inflorescence shoot apex that includes the shoot apical meristem producing floral meristem. In E6L1-OX, overexpressed *E6L1* may stimulate the shoot apical meristem to produce more flowers than WT plants, resulting in an increase of seed pod numbers.

**Fig. 7.**
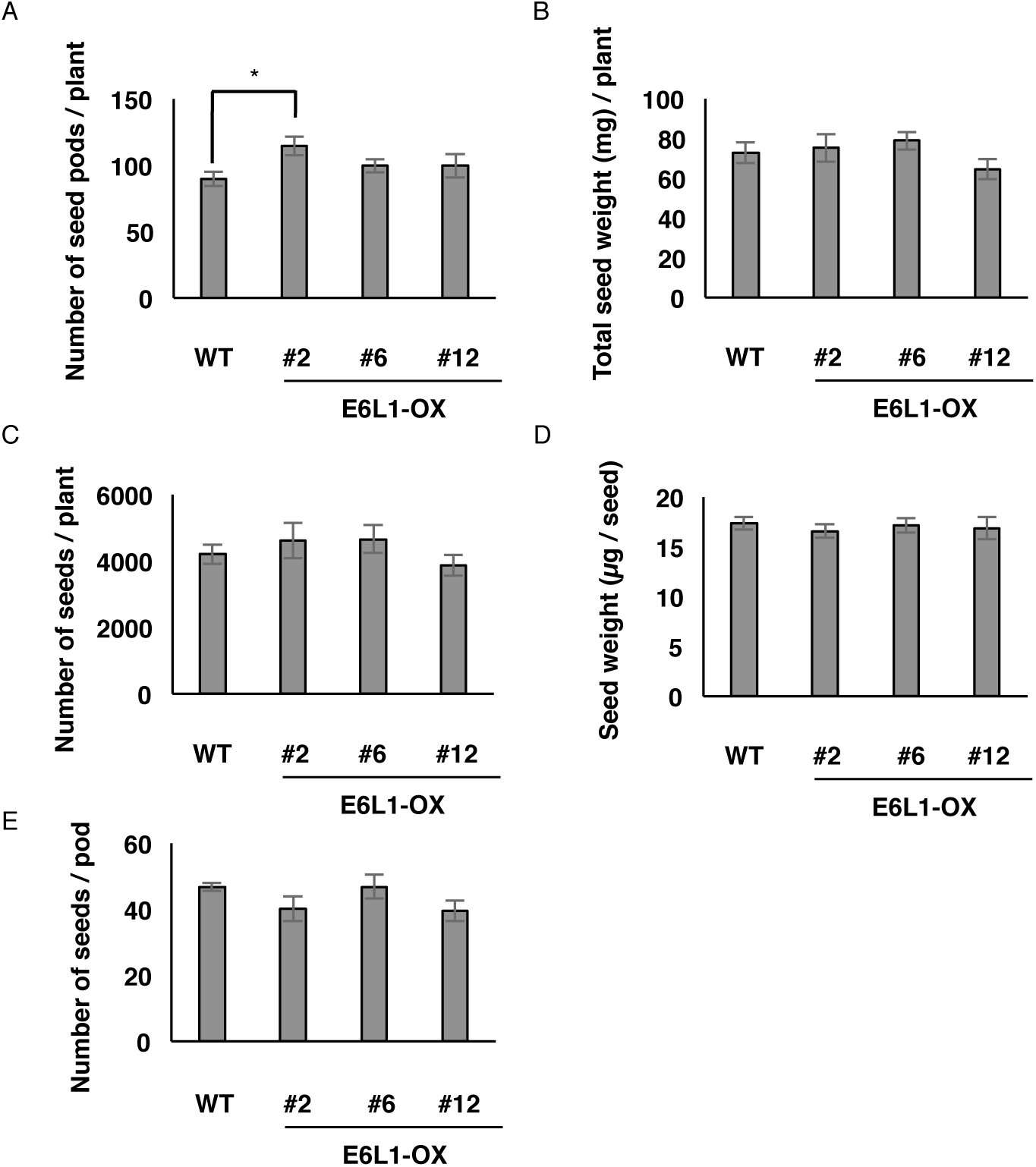
Overexpression of *E6L1* enhances seed pod number in Arabidopsis. (A) RT-PCR of *E6L1* and *EF1a* (control) transcripts in the wild type (WT) and E6L1-OX lines. (B) Number of seed pods per plant. (C) Number of seeds per plant. (D) Total weight of seeds per plant. (E) Weight per seed. (F) Number of seeds per pod. Data are means ± SEMs (*n* = 5–6 plants; * *P* < 0.05, Student’s *t* test).

## Conclusions

The results of this study showed that GmFAST, combined with *Agrobacterium*-mediated transformation using cotyledonary nodes, is an efficient and powerful method to establish homozygous soybean lines. Compared with the conventional method, which requires a large space to grow soybean plants for the selection of homozygous transgenic plants and for segregation analyses, the GmFAST method simply requires the selection of the most strongly fluorescent T2 seeds based on the expression of a fluorescent marker using a fluorescence stereomicroscope. This reduces the space requirement to at least one-tenth of that required for conventional methods (Table 1). We also generated a transgenic line with an increased seed yield. This line could be valuable in agriculture and industry. Thus, the space-saving GmFAST method will facilitate soybean transformation, which will increase the chance of developing useful soybean lines.

**Table 1.** Comparison between GmFAST and the conventional transgenic soybean selection method.

| Factors | GmFAST method | Conventional method |
| --- | --- | --- |
| Transformant identification manner | GFP fluorescence of seeds | Drug resistance of seedlings |
| Homozygote identification timing | T2 generation (seeds) | T3 generation (seedlings) |
| Segregation analysis treatment | Not required (in principle) | Required (sowing seeds in plug trays for drug selection) |
| Relative cultivation area | 0.1 | 1.0 |

## Acknowledgments

We are grateful to Makoto Hayashi (Nagahama Institute of Bio-Science and Technology, Japan) for his kind donation of the anti-OLE1 antibody, and to Masatake Kanai (National Institute for Basic Biology, Japan) and Yu Tanaka (Kyoto University, Japan) for their technical advice for plant cultivation. This work was supported; by Specially Promoted Research of Grants-in-Aid to I.H.-N. (no. 22000014), by Grants-in-Aid for Scientific Research to I.H.-N. (no. 15H05776), to T.L.S. (nos. 16K18834 and 19K05809), and to K.I. (nos. 17K15145 and 18H05496) from the Japan Society for the Promotion of Science (JSPS), by Core Research for Evolutional Science & Technology to I.H.-N. and to T.Y. (CREST, 109178) of Japan Science and Technology Agency (JST, https://www.jst.go.jp/EN/index.html), and by the Hirao Taro Foundation of KONAN GAKUEN for Academic Research to I.H.-N.

## Author contributions

K.I., T.L.S., and I.H.-N. conceived the study. K.I. and T.Y. performed experiments with *Glycine max*. TLS performed experiments with *Arabidopsis thaliana*. K.I., T.Y., T.L.S., and I.H.-N. analyzed whole data and wrote the manuscript.

## Additional information

Supplementary information is available online.

## References

Abell BM, Holbrook LA, Abenes M, Murphy DJ, Hills MJ, Moloney MM (1997) Role of the proline knot motif in oleosin endoplasmic reticulum topology and oil body targeting. Plant Cell 9:1481–1493

Bechtold N, Pelletier G (1998) In planta Agrobacterium-mediated transformation of adult Arabidopsis thaliana plants by vacuum infiltration. Methods Mol Biol 82:259–266

Clemente TE, Cahoon EB (2009) Soybean oil: genetic approaches for modification of functionality and total content. Plant Physiol 151:1030–1040

Curtis MD, Grossniklaus U (2003) A gateway cloning vector set for high-throughput functional analysis of genes in planta. Plant Physiol 133:462–469

Fischer RL, Goldberg RB (1982) Structure and flanking regions of soybean seed protein genes. Cell 29:651-660

Herman EM, Larkins BA (1999) Protein storage bodies and vacuoles. Plant Cell 11:601–614

Hu R, Fan C, Li H, Zhang Q, Fu YF (2009a) Evaluation of putative reference genes for gene expression normalization in soybean by quantitative real-time RT-PCR. BMC Mol Biol 10:93

Hu Z, Wang X, Zhan G, Liu G, Hua W, Wang H (2009b) Unusually large oilbodies are highly correlated with lower oil content in Brassica napus. Plant Cell Rep 28:541–549

Karimi M, Inze D, Depicker A (2002) GATEWAY vectors for Agrobacterium-mediated plant transformation. Trends Plant Sci 7:193–195

Kim HU, Hsieh K, Ratnayake C, Huang AH (2002) A novel group of oleosins is present inside the pollen of Arabidopsis. J Biol Chem 277:22677–22684

Lardizabal K, Effertz R, Levering C, Mai J, Pedroso MC, Jury T, Aasen E, Gruys K, Bennett K (2008) Expression of Umbelopsis ramanniana DGAT2A in seed increases oil in soybean. Plant Physiol 148:89–96

Liu WX, Liu HL, Qu le Q (2013) Embryo-specific expression of soybean oleosin altered oil body morphogenesis and increased lipid content in transgenic rice seeds. Theor Appl Genet 126:2289–2297

Meinke DW, Chen J, Beachy RN (1981) Expression of storage-protein genes during soybean seed development. Planta 153:130–139

Nakano M, Yamada T, Masuda Y, Sato Y, Kobayashi H, Ueda H, Morita R, Nishimura M, Kitamura K, Kusaba M (2014) A green-cotyledon/stay-green mutant exemplifies the ancient whole-genome duplications in soybean. Plant Cell Physiol 55:1763–1771

Nielsen NC, Dickinson CD, Cho TJ, Thanh VH, Scallon BJ, Fischer RL, Sims TL, Drews GN, Goldberg RB (1989) Characterization of the glycinin gene family in soybean. Plant Cell 1:313–328

O’Malley RC, Alonso JM, Kim CJ, Leisse TJ, Ecker JR (2007) An adapter ligation-mediated PCR method for high-throughput mapping of T-DNA inserts in the Arabidopsis genome. Nat Protoc 2:2910–2917

Rao SS, Hildebrand D (2009) Changes in oil content of transgenic soybeans expressing the yeast SLC1 gene. Lipids 44:945–951

Scallon B, Thanh VH, Floener LA, Nielsen NC (1985) Identification and characterization of DNA clones encoding group-II glycinin subunits. Theor Appl Genet 70:510–519

Schmidt MA, Barbazuk WB, Sandford M, May G, Song Z, Zhou W, Nikolau BJ, Herman EM (2011) Silencing of soybean seed storage proteins results in a rebalanced protein composition preserving seed protein content without major collateral changes in the metabolome and transcriptome. Plant Physiol 156:330–345

Schmidt MA, Herman EM (2008) Suppression of soybean oleosin produces micro-oil bodies that aggregate into oil body/ER complexes. Mol Plant 1:910–924

Shimada T, Ogawa Y, Shimada T, Hara-Nishimura I (2011) A non-destructive screenable marker, OsFAST, for identifying transgenic rice seeds. Plant Signal Behav 6:1454–1456

Shimada TL, Hara-Nishimura I (2010) Oil-body-membrane proteins and their physiological functions in plants. Biol Pharm Bull 33:360–363

Shimada TL, Shimada T, Hara-Nishimura I (2010) A rapid and non-destructive screenable marker, FAST, for identifying transformed seeds of Arabidopsis thaliana. Plant J 61:519–528

Shimada TL, Shimada T, Takahashi H, Fukao Y, Hara-Nishimura I (2008) A novel role for oleosins in freezing tolerance of oilseeds in Arabidopsis thaliana. Plant J 55:798–809

Siloto RM, Findlay K, Lopez-Villalobos A, Yeung EC, Nykiforuk CL, Moloney MM (2006) The accumulation of oleosins determines the size of seed oilbodies in Arabidopsis. Plant Cell 18:1961–1974

Tidke SA, Ramakrishna D, Kiran S, Kosturkova G, Ravishankar GA (2015) Nutraceutical Potential of Soybean: Review. Asian J Clin Nutr 7:22–32

Yamada T, Mori Y, Yasue K, Maruyama N, Kitamura K, Abe J (2014) Knockdown of the 7S globulin subunits shifts distribution of nitrogen sources to the residual protein fraction in transgenic soybean seeds. Plant Cell Rep 33:1963–1976

Yamada T, Watanabe S, Arai M, Harada K, Kitamura K (2010) Cotyledonary node pre-wounding with a micro-brush increased frequency of *Agrobacterium*-mediated transformation in soybean. Plant Biotechnology 27:217–220

