## Supplemental Figure 1 for "A space-saving visual screening method, *Glycine max* FAST, for generating transgenic soybean"

### Electronic supplementary materials

**Title:**



**Journal:**

*Journal of Plant Research*



**Corresponding author:**

Ikuko Hara-Nishimura

Faculty of Science and Engineering, Konan University, Kobe 658-8501, Japan

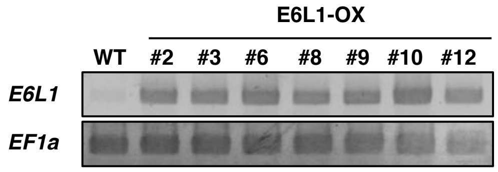


**Fig. S1 RT-PCR of *E6L1* and in *EF1a* (control) transcripts in the wild type (WT) and E6L1-OX lines.**
